# How body size and microclimate determine when solar absorptivity matters for insect body temperature

**DOI:** 10.64898/2026.10.25.761750

**Authors:** Avichal Tatu, Michael Kearney, Laura Ospina-Rozo, Nanfang Yu, Ann Roberts, Devi Stuart-Fox

**Affiliations:** School of BioSciences, The University of Melbourne, Parkville, Victoria 3010, Australia; Department of Applied Physics and Applied Mathematics, Columbia University, New York, NY 10027, USA; School of Physics, The University of Melbourne, Parkville, Victoria 3010, Australia

**Keywords:** beetles, biophysical modelling, body size, body temperature, ectotherm, microclimate, solar absorptivity, thermal melanism

## Abstract

1. The net effect of solar absorption on body temperature is difficult to intuit because it depends on nonlinear interactions between the traits of organisms and their immediate environment.
2. We used biophysical modelling to quantify solar absorption effects for Australian jewel beetles across their wide natural range of solar absorptivity (0.67-0.93), mass (more than 2 orders of magnitude) and shape under naturally co-occurring combinations of microclimatic conditions across Australia.
3. Under hot, calm conditions, solar absorptivity could increase steady-state body temperature by 2.3°C on average. Higher wind reduced this effect by more than half, whereas increasing shortwave radiation amplified it. Transient simulations indicated substantial damping of these effects at a time scale of one minute.
4. Across daylight hours during the spring–early summer period when adult jewel beetles are active, the absorptivity effect averaged 0.36°C across Australia. Under locally extreme high-radiation, low-wind conditions, the effect reached 5.69°C in the largest beetles and was more than twice as large at maximum as at minimum body mass.
5. Whether solar absorptivity is of biological significance to heat budgets is therefore highly contextual. Experiments to infer the adaptive significance of insect colour for heat budgets should account for transient responses under realistic environmental combinations.

## Introduction

The body temperature of an ectotherm governs its ability to move, feed, escape from predators and reproduce (Angilletta et al., 2002; Kingsolver, 2009). Traits that affect body temperature are therefore expected to be under strong selection. Yet the magnitude of the effect of different traits on body temperature – and therefore the strength of selection – can be difficult to predict because it depends on complex interactions between different organismal traits and surrounding environmental conditions (Porter & Gates, 1969; Kearney & Porter, 2009). The effects of traits such as colour and size on body temperature are well studied (Willmer & Unwin, 1981; Clusella Trullas et al., 2007). However, these traits are often considered in isolation or under a limited set of conditions. Few studies have systematically quantified the nature and magnitude of thermal benefits across a range of interacting trait values, or across the breadth of environmental space. This is essential if we are to understand the evolution of thermally relevant traits under current and future environmental conditions.

Colour is a trait that intuition says will be of great importance to body temperature because, all else being equal, darker surfaces absorb more solar radiation than light surfaces. Thus the thermal melanism hypothesis predicts that the warming effect of darker pigmentation would favour any genes associated with melanism in cold environments (Clusella Trullas et al., 2007; Clusella-Trullas et al., 2008). Here, colour is a proxy for absorptivity of solar (shortwave) radiation, which spans both visible and near-infrared (NIR) wavelengths. However, colour can be a poor proxy; two organisms that appear identical to the human eye may differ substantially in heat gain if their NIR absorptivity differs (Ospina-Rozo et al., 2022; Stuart-Fox et al., 2017; Wang et al., 2021). Another trait influencing how radiation affects the heat budget is the emissivity of thermal (longwave) radiation. For living organisms, emissivity is often assumed to lie between 0.90 and 0.98 due to the high water content of biological tissues (McCafferty et al., 2011; Eastick et al., 2019; Pincebourde et al., 2021). However, there is increasing recognition that emissivity can vary more than previously assumed and can strongly affect body temperature (Krishna et al., 2020). Optical traits (solar absorptivity and emissivity) interact with other morphological traits such as body size and shape. Body shape can modify many heat-transfer processes because geometry changes projected area, total surface area and characteristic dimension, and therefore both radiative and convective heat exchange (Mitchell, 1976; Kingsolver & Moffat, 1982). At a given body mass, differences in shape can alter the balance between heat gain and heat loss independently of size or absorptivity (Mitchell, 1976; Kingsolver & Moffat, 1982). Thus, the thermal effect of any one morphological trait can depend on the values of other traits that influence the same heat-transfer processes.

The magnitude of the effect of morphological traits can vary substantially depending on microclimatic conditions, making it difficult to generalise the effect across different environments. For example, solar radiation determines the shortwave energy available for absorption, and air temperature and wind influence how readily that absorbed heat is exchanged through convection (Digby, 1955; Porter & Gates, 1969; Stevenson, 1985). One way to estimate how the magnitude of the absorptivity effect changes across environments is to vary air temperature, radiation and wind independently across their ranges (e.g. de Jong et al., 1996). However, these variables do not occur in arbitrary combinations in nature. The combinations of radiation, air temperature and wind depend on location, season, time of day and microhabitat, producing a constrained environmental space (Porter & Gates, 1969).

Treating each variable independently can therefore overrepresent combinations that are rare or generate combinations that rarely or never occur in nature. To estimate the magnitude of effects of morphological traits on body temperature, it is important to consider real microclimatic conditions that preserve patterns of covariation among environmental variables.

Biophysical models of heat exchange are a powerful approach to evaluate the interactive effects of organismal traits and environmental variables on body temperature. These models calculate the heat budget of an organism, set by the balance of radiative, convective and conductive heat fluxes between the organism and its environment, as a function of organismal traits and environmental inputs (Porter & Gates, 1969; Kearney & Porter, 2009). The models allow traits such as absorptivity, body size and shape to be varied independently, while environmental conditions can be sampled from realised microclimate data that retain the relationships among radiation, air temperature and wind. This makes it possible to quantify the relative effects of individual traits and determine how their importance changes across the conditions insects experience.

Here, we use Australian jewel beetles (Buprestidae) to quantify how solar absorptivity contributes to body temperature relative to body size and shape, and how these effects change across environmental conditions. The importance of solar absorptivity for temperature regulation in insects is debated; studies of insect colour have found effects ranging from negligible (Hadley, 1970; Veron, 1974; Umbers et al., 2013) to substantial differences of up to 8.3°C (Joern, 1981). Small ectotherms such as insects are tightly coupled to their local microclimate, while body size influences both heat exchange and thermal inertia (Stevenson, 1985; Kearney et al., 2021; Pincebourde et al., 2021). The same difference in absorptivity may therefore produce a smaller temperature difference in a small insect than in a larger one (Willmer & Unwin, 1981). It is therefore important to understand the body sizes and environmental conditions under which solar absorptivity – or colour – has a substantial effect on insect body temperature.

Jewel beetles are a useful system for evaluating the thermal effect of solar absorptivity because they are diurnally active, arboreal insects that perch and forage on flowering vegetation in direct sunlight (Bellamy et al., 2013; pers. obs.), where radiative and convective heat exchange should be important. They vary substantially in body mass and solar absorptivity across both visible and NIR wavelengths, and NIR absorptivity has previously been shown to influence heating rates in jewel beetles (Wang et al., 2021). We tested model predictions against field measurements and then used empirically measured trait ranges in local and global sensitivity analyses to quantify the effects and relative importance of absorptivity, body mass and shape for steady-state body temperature, body temperature after 60 s, and time to equilibrium. We tested whether the thermal effect of absorptivity changed with body size and mapped its magnitude across Australia during the active period of the beetles. We therefore asked when absorptivity has a large effect on body temperature, how that effect changes with body size and shape, and whether the traits that matter most differ between transient and steady-state body temperatures.

## Methods

### Replication Statement

**Table 1.** Replication statement. No experimental treatments were applied; sample sizes are reported at the scale used to define empirical trait ranges and validate model predictions.

| Scale of inference | Scale at which factor is applied | Number of replicates at the appropriate scale |
| --- | --- | --- |
| Variation in body mass and shape among Australian jewel beetles | Individual beetle | 90 individuals across 18 species |
| Variation in solar absorptivity among jewel beetles | Individual beetle/species | 4 newly measured species (5 individuals per species), combined with published data for 17 species; 20 species total after overlap |
| Variation in thermal emissivity among jewel beetles | Individual beetle | 25 individuals across 5 species (5 individuals per species) |
| Field validation of predicted body temperature | Individual beetle observation | Victoria: 58 observations (50 with matched local environmental measurements); Western Australia: 31 observations |
| Field validation of operative temperature | Individual perch | Victoria: 58 observations (50 with matched local environmental measurements); Western Australia: 30 observations |

### Biophysical model

We modelled beetle body temperature using a biophysical heat-budget model implemented in NicheMapR (Kearney & Porter, 2020). For each combination of organismal traits and environmental conditions, the model calculates radiative, convective and conductive heat fluxes and solves for the steady-state body temperature (*T_c_*_,*final*_) at which heat gains and losses balance. It also calculates the thermal time constant (*τ*), which depends on the thermal capacitance of the body and the rate of heat exchange with the environment. These model outputs were then used to calculate the change in body temperature through time under constant environmental conditions:

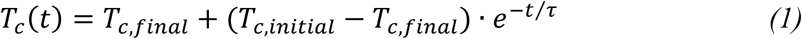

where *T_c_*(*t*)is body temperature at time *t*, *T_c_*_,*final*_ is the final steady-state body temperature, *T_c_*_,*initial*_ is the initial body temperature, and *τ* is the thermal time constant governing the rate at which body temperature approaches *T_c_*_,*final*_. From eq. 1, the beetle completes 63% of its adjustment toward *T_c_*_,*final*_ after 1τ, 95% after 3τ, and 99.3% after 5τ. We therefore define time to equilibrium as:

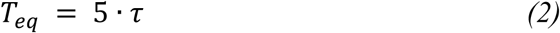

For each simulation, we extracted three response variables: steady-state body temperature (*T_c_*_,*final*_), body temperature after 60 s (*T_c_*_,60_), and time to equilibrium (*T_eq_*).

Transient simulations were initialised with body temperature equal to air temperature, and *T_c_*_,60_ was extracted after 60 s. We used 60 s as a standardised short-term interval to represent brief perch-scale exposure as beetles move among microhabitats, and to compare transient thermal responses with steady-state predictions.

To isolate the effects of the focal morphological traits and environmental conditions, posture and substrate properties were held constant, and metabolic heat production was set to zero. Fixed model parameters and settings are provided in Supporting Information Table S1.

### Model parameterisation

We parameterised the model using empirical measurements of optical and morphological traits from jewel beetles. Traits were measured to define biologically realistic bounds for sensitivity analyses rather than to compare species as fixed categories. Sample sizes and scales of inference for trait measurements and field validation are summarised in the replication statement (Table 1).

### Solar absorptivity (α)

We combined published reflectance data for 17 jewel beetle species (Wang et al., 2021) with measurements from four additional species. Reflectance was characterised across 300–1700 nm and weighted by the incident solar spectrum to calculate broadband solar reflectivity, from which solar absorptivity (*α*) was then calculated as 1 − *R*, where *R* is broadband solar reflectivity expressed as a proportion. The combined dataset comprised 20 species and yielded solar absorptivities of 0.67–0.93, which were used as the empirical bounds for subsequent analyses (Supporting Information Methods S1).

### Mid-infrared emissivity (ε)

We measured mid-infrared spectral reflectance and transmittance across 3–16 µm using Fourier-transform infrared spectroscopy (FTIR) in five jewel beetle species. Spectral emissivity was then calculated as

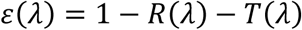

following Kirchhoff’s law (Tsai et al., 2020), and broadband emissivity was obtained by weighting spectral emissivity by the Planck distribution across biologically relevant temperatures. Broadband emissivity varied little among measured individuals and species (0.88–0.90), so we fixed emissivity at the median value (*ε* = 0.90) in all subsequent analyses (Supporting Information Methods S2).

### Shape and mass

Morphological measurements were taken from 90 individuals across 18 jewel beetle species selected to span a broad range of body sizes and masses. We approximated beetle body shape as a tri-axial ellipsoid characterised by maximum body length (a), width (b) and height (c). Relative width (b/a) and relative height (c/a) were used as descriptors of body proportions allowing variation in body shape to be separated from overall body size. In the model, body mass determined body volume, while relative width and relative height determined how this volume was distributed among the three ellipsoid axes (Supporting Information Methods S3).

### Field measurements for validation

To validate the model across contrasting thermal environments, we collected beetle body temperatures, operative temperatures and local environmental measurements for *Temognatha bonvouloirii* in Western Australia and *Castiarina sexplagiata* in Victoria. Air temperatures associated with beetle observations ranged from 24.8 to 47.4°C in Western Australia and from 17.8 to 34.2°C in Victoria. Sampling periods, sample sizes and regional microclimate reconstruction settings are provided in Supporting Information Table S2.

Beetle body temperature was measured non-invasively using infrared thermography, with emissivity set to 0.90 based on our measurements. Body temperature was extracted from the dorsal surface. Surface temperature was used as a proxy for body temperature because the calculated Biot number was <0.1 (Porter et al., 1973), indicating negligible internal thermal gradients.

Local air temperature, wind speed, ground temperature and sky temperature were measured during field sampling and matched to beetle observations using the corresponding environmental records (Supporting Information Methods S4).

### Operative temperature (Te)

We estimated operative temperature using hollow copper beetle models cast from moulds of dead beetles so that model size and external shape matched the focal species. The copper surface was sanded to produce a dull finish. Each model contained an internal Type K thermocouple (RS PRO, 0.2 mm diameter) connected to a four-channel wireless datalogger thermometer (CENTER 521; Center Technology Corp., Taiwan). Copper was used because its high thermal conductivity allows rapid equilibration with local environmental conditions.

Operative temperature is the steady-state temperature of an inanimate model exposed to the same microclimate (Bakken, 1992; Dzialowski, 2005). As beetles flew frequently between perches, an individual observed at a given location may have arrived only moments earlier and retained a body temperature from its previous microclimate rather than the immediate local conditions. We therefore used operative temperature to represent the thermal conditions at each perch and to evaluate the steady-state model predictions.

Preliminary observations showed that copper-model temperature stabilised within 30 s. Whenever a beetle was observed, a copper model was therefore placed in the same location for 30 s, after which temperature was recorded.

### Model validation

We validated model predictions using two complementary approaches. In the first, the model was driven by hourly environmental conditions reconstructed using micro_silo, the NicheMapR interface that uses Australian SILO climate data to drive the NicheMapR microclimate model (Kearney & Porter, 2017; Jeffrey et al., 2001). Environmental conditions were reconstructed for each field region during the relevant sampling windows. For Victoria, micro_silo was parameterised at a representative Cardinia coordinate (37.9678°S, 145.3987°E), with a slope of 35°, aspect of 0° and a wind factor of 0.5. For Western Australia, it was parameterised at a representative coordinate (29.5785°S, 117.0477°E), with no slope or aspect correction, and a wind factor of 1.0. Hourly air temperature, wind speed, shortwave radiation, solar zenith angle, sky temperature and ground temperature were used as inputs to the biophysical model of the beetle.

In the second approach, the model was driven primarily by local environmental measurements collected in the field. Air temperature, wind speed, ground temperature and sky temperature were obtained from the field measurements, with radiative temperature calculated as the mean of measured ground and sky temperature. Shortwave radiation and solar zenith angle were not measured in the field and were therefore obtained from the nearest *micro_silo* timestamp within 60 min.

Observations were matched to predictions using the nearest timestamp within 60 min. Predicted *T_c_*_,*final*_ was compared with observed beetle body temperature and operative temperature. Model performance was evaluated using root mean square error (RMSE), bias, and Pearson’s correlation coefficient (r).

### Climate blocks

To assess how morphological traits influence body temperature across environmental conditions, we used precalculated hourly microclimate for Australia from the MicroclimOz data set (Kearney, 2019). This dataset includes air temperature, wind speed, solar radiation, solar zenith angle, sky temperature and ground temperature from 1990 to 2017, simulated at 5km resolution over ∼2000 points. We sampled across the full 28-year record to capture a broad range of conditions occurring across Australia rather than conditions from a single year. We created six blocks representing hot daytime, cool daytime and cold nighttime conditions. Each was divided into calm and windy states. Daylight was defined as solar zenith angle < 90° and night as solar zenith angle ≥ 90°. Cool daytime conditions were restricted to hours at or before solar noon. Solar noon was identified as the hour of minimum solar zenith angle for each grid cell and day.

Wind conditions were defined separately for day and night because the two periods have different wind distributions. Windy conditions were defined using the 75th percentile of wind speed within the daytime or nighttime period. Calm daytime conditions were defined as wind speed ≤ 1.0 m s⁻¹. This gave a simple absolute threshold for low convective cooling during solar exposure. At night there is no shortwave heating, so we instead used the 10th percentile of nighttime wind speed to represent the calmest part of the nighttime distribution. Hot daytime hours were selected from the upper tail of daytime air temperature. We started at the 99th percentile and relaxed the threshold as needed until enough records were available. Cool daytime and cold nighttime conditions were defined using the 10th percentile of air temperature within their respective periods. We randomly sampled 60 hourly environments from each block. These blocks were designed to represent realised combinations of environmental conditions rather than matched one-variable contrasts; direct effects of individual environmental drivers were quantified separately in the one-factor-at-a-time analysis. Final thresholds used to delineate the climate blocks are provided in Supporting Information Table S3. These blocks were used in the local and global sensitivity analyses (see Sensitivity analysis).

### Sensitivity analysis

We used both local and global sensitivity analyses because they address different aspects of parameter importance. The local sensitivity analysis tested how model output changed when one parameter was varied at a time. The global sensitivity analysis tested the relative contribution of each parameter when all parameters were varied together, including their interactions.

For local sensitivity analysis, we used a one-factor-at-a-time approach. Within each sampled hourly environment, we varied one focal trait across its empirical range while holding all other traits at the midpoints of their observed ranges. This was done for solar absorptivity, body mass, and the two ellipsoidal shape parameters (relative width and relative height), and the model was rerun to estimate steady-state body temperature (*T_c_*_,*final*_), transient temperature at 60 s (*T_c_*_,60_), and time to equilibrium (*T_eq_*). Trait effects were quantified as the difference in model output between the maximum and minimum value of the focal trait. We also calculated an absorptivity × mass interaction term as the difference in the absorptivity effect between maximum and minimum body mass.

We applied the same one-factor-at-a-time analysis to environmental drivers. We varied air temperature, shortwave radiation, wind speed, sky temperature, ground temperature and solar zenith angle across the 5th–95th percentile range represented in the combined sampled climate panel. For these analyses, the remaining environmental variables retained the values of each sampled environment. This allowed us to compare the effects of changing environmental conditions with the effects of traits.

To quantify how environmental conditions altered the thermal effect of each trait, we varied each environmental driver across the same 5th–95th percentile range while calculating the difference in model output between the maximum and minimum value of each focal trait, with all other environmental variables held at their overall climate-panel medians. Because environmental variables were varied one at a time, these simulations did not retain the covariance among environmental variables.

For global sensitivity analysis, we used Sobol variance decomposition implemented in the R package *sensitivity* (Iooss et al., 2024). Traits were sampled independently and uniformly across their observed ranges. We did this to separate the effects of individual traits rather than preserve the trait covariance found in natural populations where body size and reflectance covary (Wang et al., 2023). The Sobol indices therefore show how sensitive the model was to each trait across these ranges, rather than the variation expected among real beetles. We estimated first-order and total-order Sobol indices using the Jansen estimator (Jansen, 1999), as implemented in the soboljansen() function. First-order indices quantify the variance attributable to each parameter alone, whereas total-order indices quantify the variance attributable to each parameter including all interaction effects.

We selected one representative environment from each climate block. For each block, environmental variables were standardised and Euclidean distance was used to select the observed environment closest to the median conditions. We used the same six environments for every Sobol sample so that differences were due to the traits rather than changes in the environment. For each sampled trait combination, we predicted *T_c_*_,*final*_, *T_c_*_,60_ and *T_eq_* across the six representative climates and averaged responses equally across blocks. Analyses used 2500 base samples and 1000 bootstrap replicates to derive 95% confidence intervals for each Sobol index.

### Spatial modelling of absorptivity effects

We quantified geographic variation in the thermal effect of absorptivity by applying the biophysical model to hourly gridded MicroclimOz rasters across Australia for October– December 2017. Australia provides a useful climate space for this analysis because of both its size and climatic diversity. The continent spans approximately 33° of latitude, from about 11°S to 44°S, equivalent to the latitudinal range from around 11°N to 44°N in the Northern Hemisphere. Australia spans a wide range of climates, from the tropical north and dry interior to temperate regions in the south and southeast. Mediterranean-type climates occur in the southwest, while alpine conditions are found in the southeastern highlands (Peel et al., 2007). This gives Australia a broad range of conditions, including environments where insect activity can be limited by both low and high temperatures.

October–December was selected as a focal spring to summer period when adult jewel beetles are commonly active above ground and exposed to solar radiation. We used 2017 because it captured most of the environmental variation occurring during the same period across the full 1990–2017 dataset. For each year, we calculated how much of the variation across the six environmental variables from the full 1990–2017 period it covered. In 2017, this was 98.1%.

The analyses used air temperature, wind speed, shortwave radiation, solar zenith angle, sky temperature and ground temperature, aligned to a common grid and masked to Australian land cells. Simulations were restricted to daylight hours which were defined as cosine of solar zenith angle > 0. For each grid cell and hour, we predicted steady-state body temperature (*T_c_*_,*final*_) at minimum and maximum absorptivity and calculated Δ*T_c_*_,*final*_ as the difference between them, thereby isolating the thermal effect of absorptivity while holding other traits at the midpoints of their observed ranges. We repeated the simulations at three levels of wind exposure: no reduction in wind speed, 70% reduction in wind speeds > 1.5 m s⁻¹, and 90% reduction in wind speeds > 1.5 m s⁻¹. All other environmental variables were left unchanged.

Using the result of the sensitivity analysis, we identified the climatically extreme conditions under which solar absorptivity was expected to have the greatest thermal leverage. For every daylight hour, we calculated the ratio of shortwave radiation to wind speed as shortwave radiation / (wind speed + 0.1), such that hours with high solar radiation and low wind received the highest scores. Then, for each grid cell, we selected the 10 hours with the highest scores and calculated mean Δ*T_c_*_,*final*_ across these hours. The constant of 0.1 prevents division by zero when wind speed approaches zero. We used the top 10 hours rather than a single hour so that the results represented the actual local extreme conditions rather than one anomalous hour. We calculated this at the minimum and maximum observed body mass using the same hours for both body sizes to examine how body size affected the thermal effect of absorptivity under extreme conditions.

## Results

### Trait variation

Solar absorptivity ranged from 0.67 to 0.93. Body mass ranged from 0.008 to 2.64 g, spanning approximately 2.5 orders of magnitude. Relative width (b/a) ranged from 0.27 to 0.63 and relative height (c/a) from 0.23 to 0.34.

### Validation

Model performance differed between sites, with lower RMSE and higher correlations when predictions were driven by measured local conditions rather than reconstructed microclimate (Fig. 2). For body temperature, reconstructed inputs gave r = 0.54 (RMSE = 5.81°C) in Victoria and r = 0.85 (RMSE = 3.34°C) in Western Australia, compared with r = 0.71 (RMSE = 3.96°C) and r = 0.91 (RMSE = 2.40°C), respectively, using local inputs. Agreement with operative temperature using local inputs was also strong in both regions (r = 0.89 and 0.92, respectively). Complete validation statistics are provided in Supporting Information Table S4.

**Figure 1.**
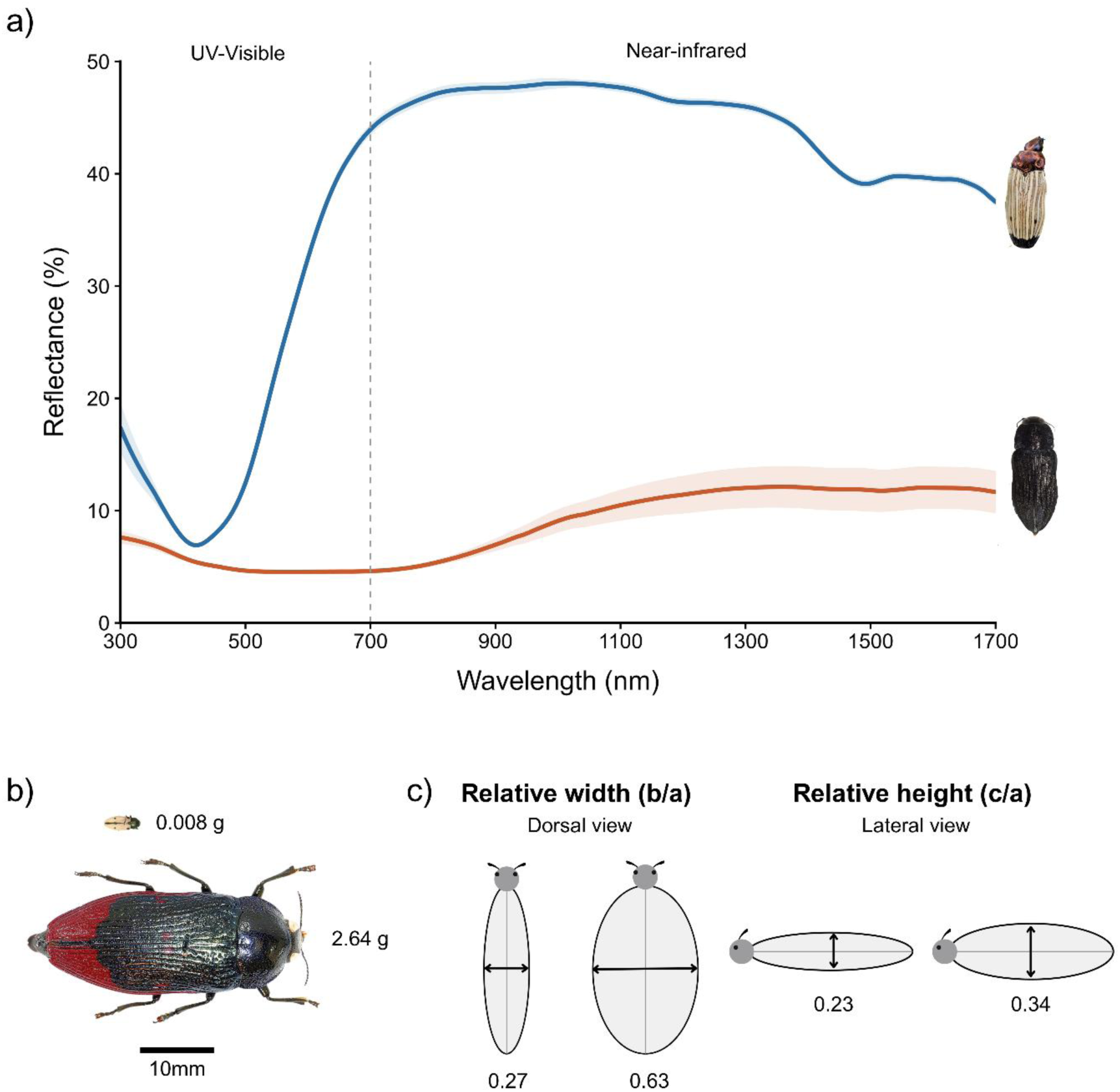
Trait variation used to parameterise the biophysical model. (a) Mean reflectance spectra (± SD) for *Castiarina luteipennis* and *Merimna atrata* from Wang et al. (2021), spanning the UV–visible (300–700 nm) and near-infrared (700–1700 nm) regions. (b) Representative body-size extremes, with *Castiarina sexguttata* (0.008 g, 6 mm) and *Temognatha bonvouloirii* (2.64 g, 44.8 mm) shown to scale. (c) Observed extremes in relative width (b/a: 0.27, *C. bella*; 0.63, *C. thomsonii*) and relative height (c/a: 0.23, *C. campestris*; 0.34, *C. thomsonii*) used to parameterise the ellipsoid geometry.

**Figure 2.**
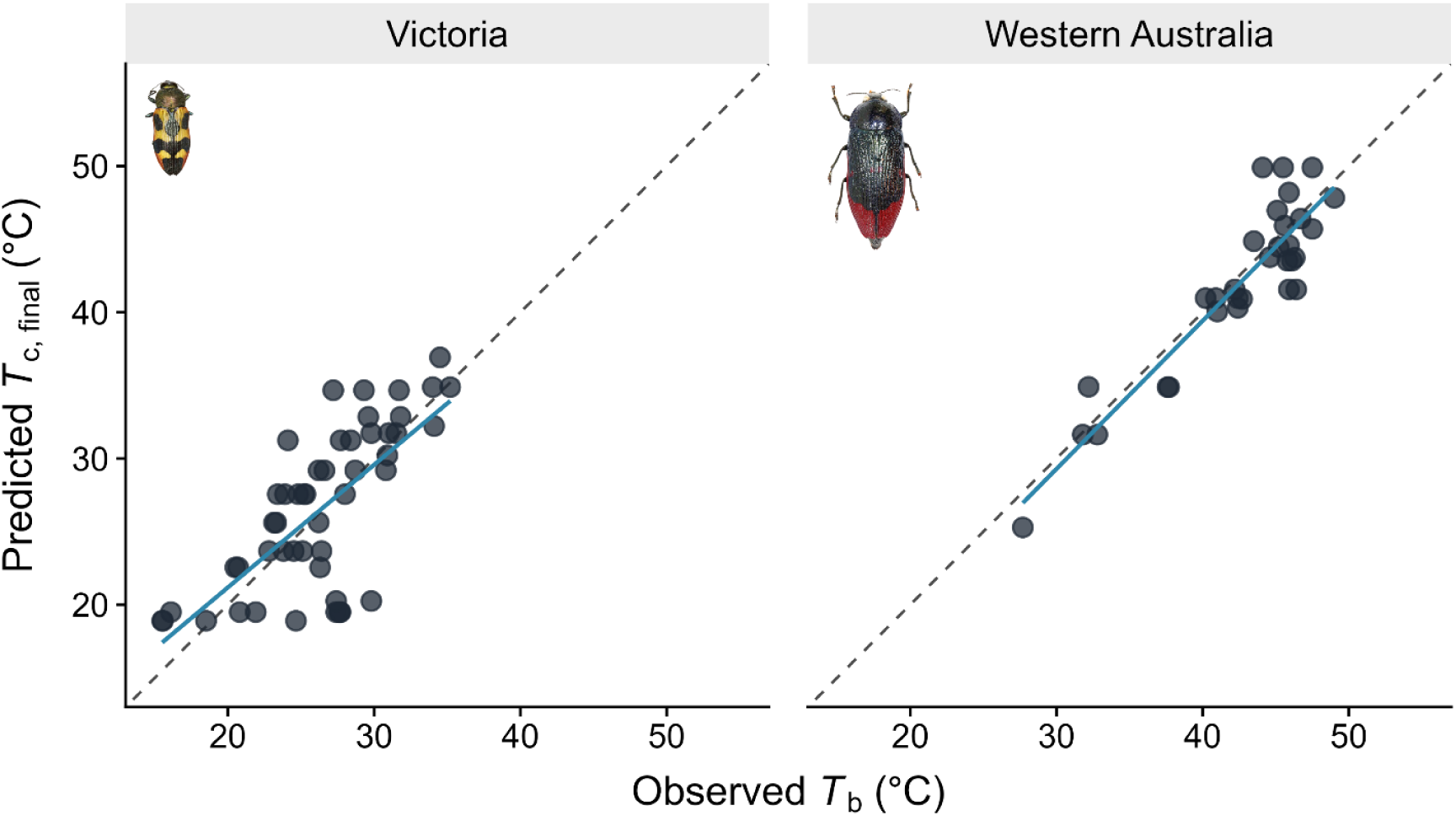
Validation of biophysical model predictions against field observations in Victoria (*Castiarina sexplagiata*) and Western Australia (*Temognatha bonvouloirii*). Predicted steady-state body temperature (*T_c_*_,*final*_), calculated using measured local environmental conditions, is plotted against observed body temperature (*T_b_*). The dashed line represents the 1:1 relationship between predicted and observed temperature, and the solid line shows the fitted linear relationship.

### Local sensitivity analysis

The effect of absorptivity depended strongly on environmental conditions. Under hot, calm conditions, increasing absorptivity across its observed range increased *T_c_*_,*final*_ by 2.3°C on average, with effects reaching 3.24°C. Under hot, windy conditions, the mean effect fell to 1.05°C, with a maximum of 1.42°C. Mean effects were smaller under cooler daytime conditions (0.37–0.93°C) and were effectively zero at night (Fig. 3).

**Figure 3.**
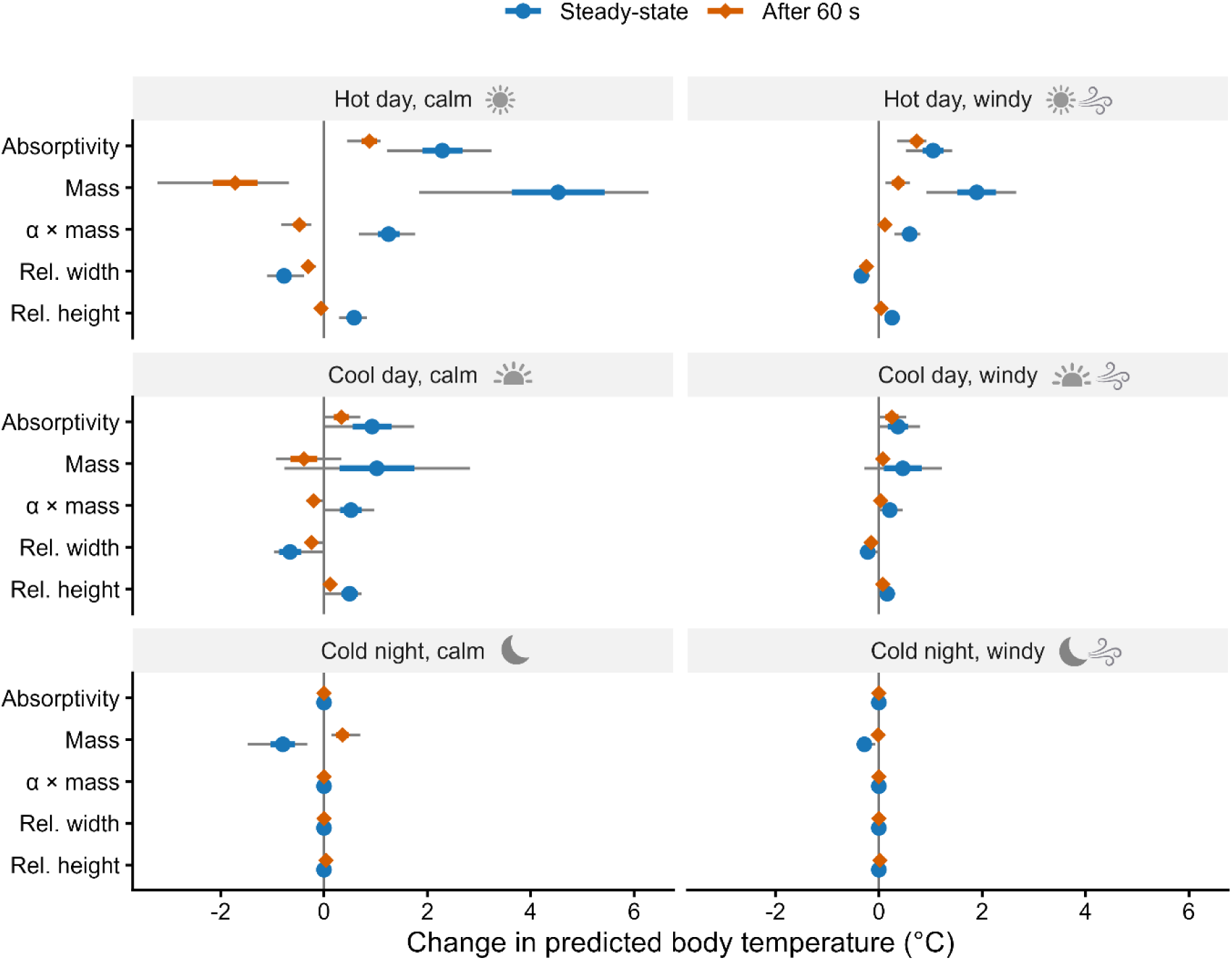
Local sensitivity of predicted body temperature to morphological traits across six climate blocks. One-factor-at-a-time effects of solar absorptivity, body mass, relative width, relative height and the absorptivity × mass interaction are shown for steady-state body temperature (Δ*T_c_*_,final_; blue circles) and body temperature after 60 s (Δ*T_c_*_,60_; orange diamonds). Climate blocks represent hot and cool daytime and cold nighttime conditions, each under calm and windy conditions. Points show the mean effect across 60 realised hourly environments within each block, coloured bars show ±1 SD and grey bars show the minimum–maximum range. Trait effects are calculated as the difference in model output between the maximum and minimum observed value of each trait. The absorptivity × mass interaction is the difference in the effect of absorptivity between maximum and minimum body mass. The vertical line at zero indicates no effect; positive values indicate that increasing the trait increases predicted body temperature, whereas negative values indicate the opposite.

The mean effect of mass on *T_c_*_,*final*_ was larger in magnitude than that of the other traits in each climate block. Across the four daytime blocks, the mean effect of increasing mass ranged from 0.46 to 4.53°C. At night, the direction reversed: increasing mass reduced *T_c_*_,*final*_ by 0.80°C on average under calm conditions and by 0.28°C under windy conditions. The absorptivity × mass interaction was strongest under hot, calm conditions, with a mean interaction effect of 1.25°C and a maximum of 1.76°C.

Body shape had smaller effects during the day. Increasing relative width reduced *T_c_*_,*final*_ by 0.21–0.78°C on average across the four daytime blocks, whereas increasing relative height increased it by 0.16–0.58°C. Neither shape variable had an appreciable effect at night.

Trait effects after 60 s were generally smaller than at steady state. Under hot, calm conditions, increasing mass reduced *T_c_*_,60_ by 1.72°C on average, whereas increasing absorptivity raised it by 0.88°C. Under hot, windy conditions, absorptivity increased *T_c_*_,60_ by 0.73°C compared with a 0.37°C increase due to mass. Effects were smaller under cooler conditions (Fig. 3). Increasing mass prolonged *T_eq_* by an average of 633 s across climate blocks. Relative height had a smaller effect, increasing it by 56 s on average, whereas relative width changed it by less than 1 s. Absorptivity had no effect on *T_eq_* (Supporting Information Fig. S2).

### Environmental effects on trait sensitivity

Environmental conditions strongly altered the steady-state effects of absorptivity and body mass. Shortwave radiation produced the largest change in the effect of absorptivity on ***T_c_***_,***final***_ (2.17°C) and strongly altered the effect of body mass (3.69°C; Supporting Information Fig. S3). Direct effects of the environmental variables on ***T_c_***_,***final***_, ***T_c_***_,**60**_, and ***T_eq_*** are shown in Supporting Information Fig. S4.

### Global sensitivity analysis

Absorptivity and mass explained most of the variation in steady-state body temperature averaged across the six representative environments (Fig. 4). For *T_c_*_,*final*_, the first-order Sobol index was 0.42 for absorptivity (95% CI = 0.38–0.46) and 0.41 for mass (0.37–0.45). Total-order indices were 0.47 and 0.43, respectively. For *T_c_*_,60_, mass had a higher first-order index (0.49, 95% CI = 0.46–0.53) than absorptivity (0.42, 0.39–0.47), although their confidence intervals overlapped. Body shape contributed less to both temperature responses. Time to equilibrium was almost entirely determined by mass (S1 = 0.99; ST ≈ 1).

**Figure 4.**
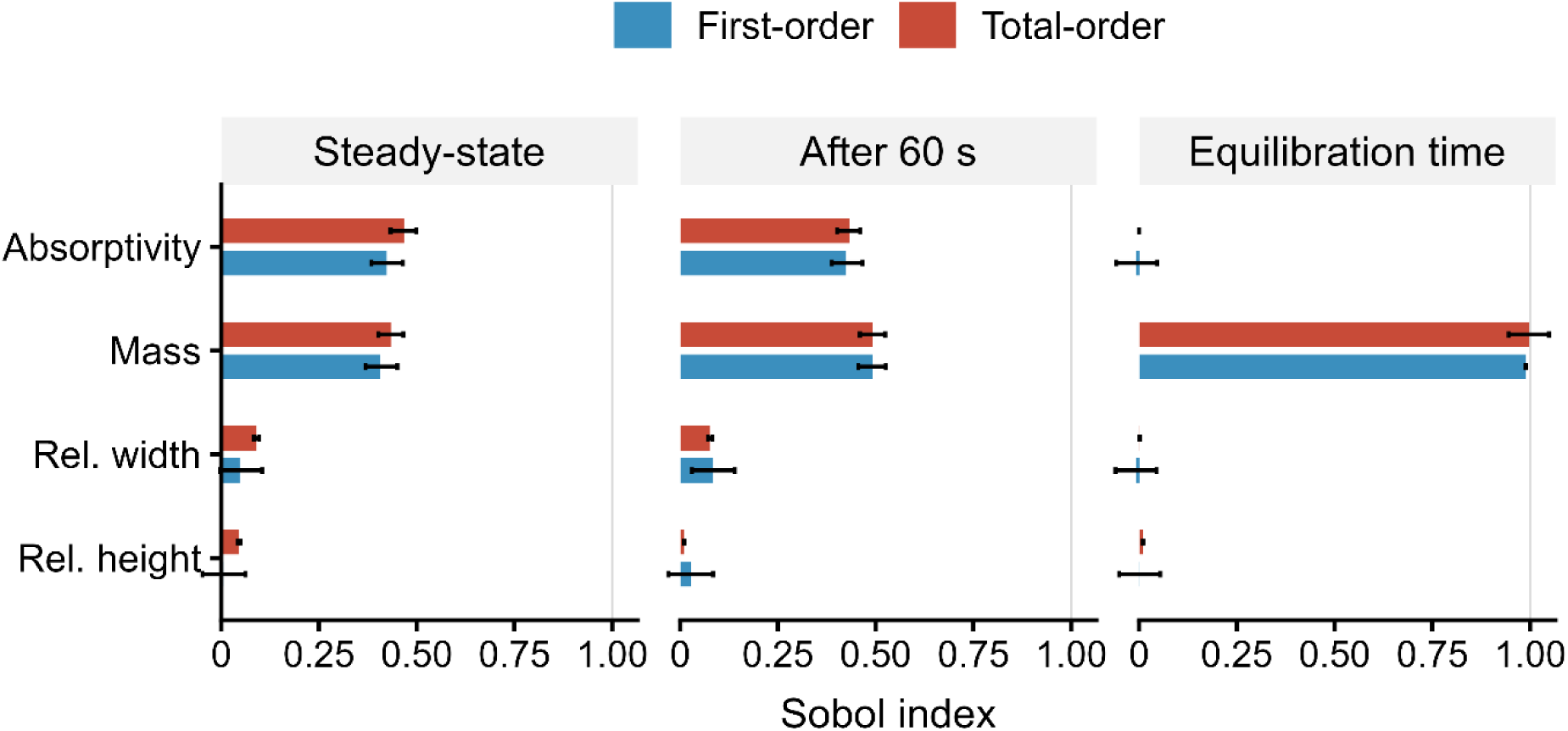
Global sensitivity of predicted thermal responses to variation in morphological traits. First-order (*S*_1_) and total-order (*S_T_*) Jansen Sobol sensitivity indices are shown for solar absorptivity, body mass, relative width and relative height across three response variables: steady-state body temperature (*T_c_*_,final_), body temperature after 60 s (*T_c_*_,60_) and time to equilibrium (*T*_eq_). First-order indices represent the contribution of each trait alone to model-output variance, whereas total-order indices include both the focal trait and its interactions with other traits. Traits were sampled independently from uniform distributions spanning their observed ranges using 2500 base samples. For each trait combination, model responses were calculated in one realised representative environment from each of the six climate blocks and averaged equally across blocks. Representative environments were the observed pixel-hours closest to the multivariate median conditions of each block. Error bars show 95% confidence intervals from 1000 bootstrap replicates.

The local and global analyses gave slightly different rankings because they measure different things. The local analysis measured the effect of changing a trait within individual environments, whereas the Sobol analysis measured variation in responses averaged across the six representative environments.

### Spatial modelling of absorptivity effects

Across daylight hours during October–December 2017, changing absorptivity across its observed range increased *T_c_*_,*final*_ by 0.36°C on average across Australia when wind was unchanged. Reducing wind speeds above 1.5 m s⁻¹ by 70% increased this to 0.53°C, while a 90% reduction increased it to 0.79°C. Spatial patterns under the three wind-exposure scenarios are provided in Supporting Information Fig. S5.

The effect was much larger during the high-radiation, low-wind hours selected for each grid cell, particularly for larger beetles. Across Australia, mean delta *T_c_*_,*final*_ during the ten highest-ranked hours was 1.21°C at minimum body mass and 2.48°C at maximum body mass. Maximum ten-hour pixel means reached 2.40°C and 5.69°C, respectively. The strongest effects for both body sizes occurred near Longreach, Queensland (Fig. 5).

**Figure 5.**
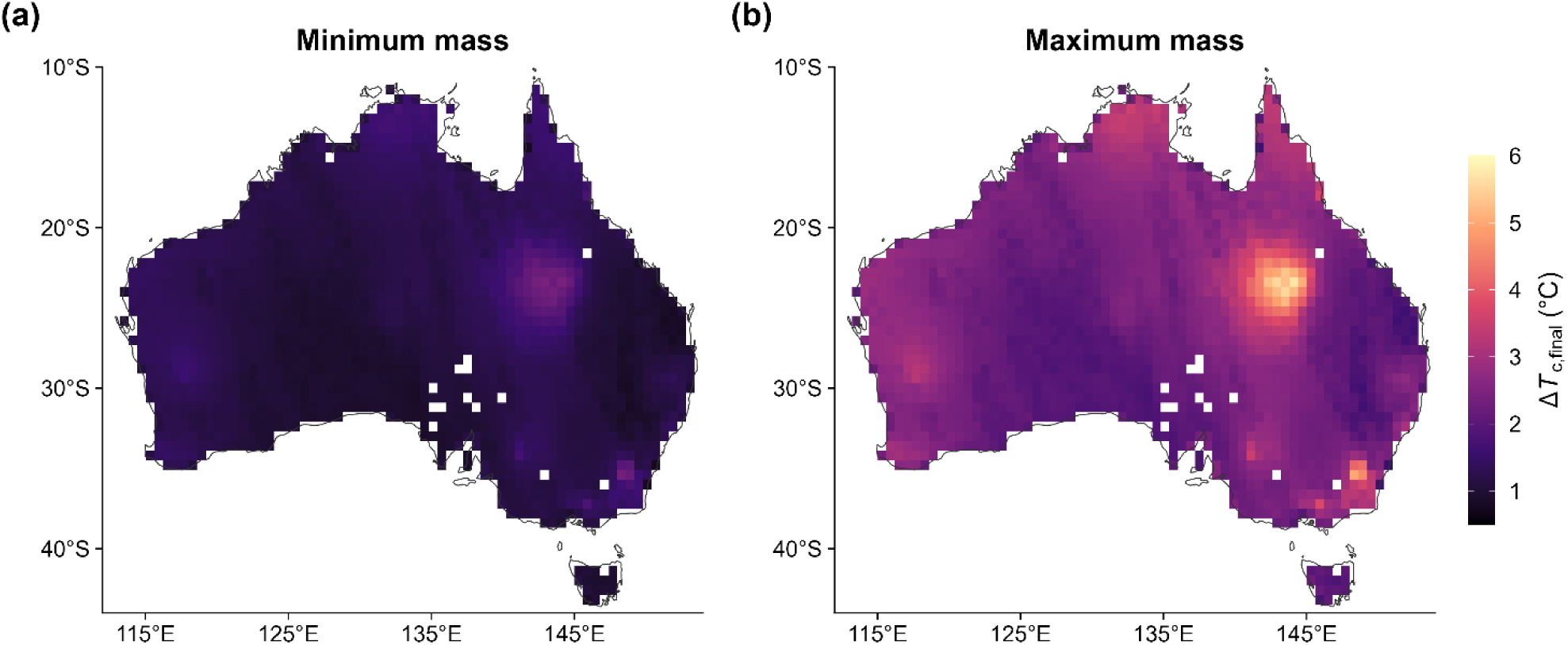
Spatial variation in the effect of solar absorptivity on predicted steady-state body temperature under locally high-radiation, low-wind conditions across Australia. For each grid cell, daylight hours during October–December 2017 were ranked according to shortwave radiation / (wind speed + 0.1 m s^-1^), and the thermal effect of absorptivity (*ΔT_c_*_,*final*_) was averaged across the ten highest-ranked hours. Δ*T_c_*_,*final*_represents the difference in predicted steady-state body temperature between the maximum (0.93) and minimum (0.67) observed solar absorptivity. Panels show predictions at (a) minimum observed body mass (0.008 g) and (b) maximum observed body mass (2.64 g), using the same selected hours for both body sizes. Relative width and relative height were held at the midpoints of their observed ranges. Higher values therefore indicate conditions in which variation in solar absorptivity produces a larger difference in steady-state body temperature.

## Discussion

We examined how much solar absorptivity contributes to beetle body temperature relative to other temperature-linked morphological traits, and how that effect varies across environmental conditions. In the global sensitivity analysis, absorptivity had the highest first-order Sobol index for steady-state body temperature, closely followed by mass, whereas mass had a higher first-order index for transient body temperature. The effect of absorptivity was strongly modulated by wind speed and body size, in addition to the expected response to solar radiation. Mass showed a different pattern. Of all the traits we considered, it produced the largest local effects across climate blocks and, as expected, had by far the strongest effect on time to equilibrium amongst all traits. Body shape affected steady-state and transient-state body temperature, although its effects were smaller overall. The spatial analysis showed the same conditional responses at a continental scale. Differences due to absorptivity were small when averaged across the beetles’ active period but reached up to 5.69°C under high-radiation and low-wind conditions. Thus, absorptivity can have a large effect on body temperature, but only under specific environmental conditions and depending on the size of the insect.

The conditional nature of the effect of absorptivity on temperature has direct consequences for how the thermal melanism hypothesis should be interpreted. The hypothesis posits that darker individuals heat faster and reach higher body temperatures and are therefore favoured in cooler environments (Clusella Trullas et al., 2007; Watt, 1968). This pattern is well-supported at the macroecological scale where lizards are darker in cooler, less exposed climates globally (Clusella-Trullas et al., 2008), butterfly and dragonfly assemblages are darker in cooler regions across Europe (Zeuss et al., 2014), and ant assemblages have darker and larger individuals in cooler environments (Bishop et al., 2016). Yet these macroecological associations do not imply a consistently large thermal effect at the scale of individual insects, with empirical estimates ranging from negligible differences under some conditions to several degrees under others (Köhler & Schielzeth, 2020; Umbers et al., 2013). Our study shows how the magnitude of the absorptivity effect changes across environmental conditions. Shortwave radiation most strongly altered the effect of absorptivity on body temperature.

This is consistent with the thermal melanism prediction that darker individuals can gain a thermal advantage in cooler environments when solar radiation is available (Clusella-Trullas et al., 2008). However, high air temperature does not necessarily mean high solar exposure. Insects can experience high temperatures in shaded microhabitats while receiving little direct solar radiation, in which case differences in absorptivity should have only a small effect on body temperature. Conversely, cooler but sun-exposed conditions can still produce a larger effect when shortwave radiation intensity is high. Wind further modifies this effect by increasing convective heat exchange and reducing the temperature difference produced by additional solar absorption (Stevenson, 1985). Importantly, these environmental variables covary in realised climates. In our climatic data, high solar radiation and high air temperature often coincided with higher wind speeds, which reduced the thermal effect of absorptivity under exposed conditions. Combinations of high radiation and low wind may therefore occur more locally where wind exposure is reduced, for example within sheltered vegetation. Air temperature alone is therefore a poor predictor of when absorptivity will have a large thermal effect.

These comparisons also depend on how absorptivity is defined. Most studies quantify colour only within the visible spectrum and use visible darkness as a proxy for solar absorptivity. Colour is typically characterised either as discrete visible morphs (de Jong et al., 1996; Köhler & Schielzeth, 2020; Watt, 1968) or as lightness extracted from digital images of specimens (Bishop et al., 2016; Zeuss et al., 2014). However, neither approach captures variation in reflectance beyond the visible spectrum. This omission is important because near-infrared wavelengths account for 55% of the radiant energy in direct sunlight (Stuart-Fox et al., 2017). In jewel beetles, heating rates are influenced more strongly by near-infrared than by visible reflectance, decreasing by approximately 0.011°C min⁻¹ for every 1% increase in near-infrared reflectivity (Wang et al., 2021). Likewise, higher absorptivity predicted faster heating and greater temperature increases after 5 min in isolated Christmas beetle elytra, when temperatures were approaching steady state (Ospina-Rozo et al., 2022). Because visible and near-infrared reflectance are only partly correlated in beetles (Wang et al., 2021, 2023), the range of absorptivity we sampled differs from the range that would be inferred from visible-spectrum measurements alone.

Emissivity showed much less variation among the jewel beetles measured here, ranging only from 0.88 to 0.90, and was therefore held constant in the sensitivity analyses. Within this system, variation in solar absorptivity was therefore a much greater source of optical variation than emissivity. However, this may not hold for taxa with a broader range of thermal emissivity.

The effect of absorptivity was not fixed but depended on beetle body size. Across the four daytime climate blocks, larger beetles reached higher mean steady-state body temperatures than smaller beetles. In the selected-hour spatial analysis, the continental mean temperature difference generated by the observed absorptivity range approximately doubled from the smallest to the largest beetles. Absorptivity therefore had greater thermal leverage over steady-state body temperature in larger beetles, as expected from theory (Stevenson, 1985; Kearney et al., 2021). Low absorptivity may consequently provide disproportionately greater protection from solar heat gain in larger beetles.

The spatial analysis showed why the magnitude of the interaction between absorptivity and body size can matter biologically. Under cool but sunny conditions in the Australian Alps and adjacent Southeastern Highlands, an air temperature of 7.6°C combined with 570 W m⁻² shortwave radiation and low wind produced steady-state body temperatures of 13.7–16.1°C across the observed absorptivity range in the smallest beetles, compared with 20.7–26.0°C in the largest (Supporting Information Table S5). Greater absorptivity could therefore provide several degrees of additional passive warming under cool conditions, potentially increasing opportunities for activity, particularly in larger beetles. The consequence was very different around Longreach-Opalton region in central-western Queensland. At the grid cells producing the largest mean absorptivity effect for each body mass, air temperatures were approximately 28°C combined with substantial solar radiation and very low wind, and increasing absorptivity raised predicted steady-state body temperature from 35.3 to 37.7°C in the smallest beetles but from 44.2 to 49.9°C in the largest (Supporting Information Table S5). Thus, the same optical difference can facilitate warming under cool conditions but add considerably to thermal load under warm, sunny and low-wind conditions, with both effects amplified by body size. This provides a mechanistic basis for the suggestion that higher NIR reflectance in larger jewel beetles may offset the thermal consequences of large body size (Wang et al., 2023).

Beyond the interactive effect with absorptivity, body size also influences how quickly body temperature responds to changing conditions. Small insects heat and cool rapidly and track changes in their local environment closely, whereas larger insects respond more slowly and retain the thermal effects of previous conditions for longer (Kearney et al., 2021; Pincebourde et al., 2021). This means that the effect of a trait can depend on when body temperature is measured. Under hot, calm conditions, increasing mass raised steady-state body temperature but lowered body temperature after 60 s. Absorptivity did not reverse direction, but its effect was much smaller after 60 s than at steady state. Because insects in the field may leave a microclimate before equilibrating, steady-state predictions may overestimate the thermal effect realised during short periods of exposure and, for mass, can even predict the opposite direction of the short-term response.

Body size also affects steady-state temperature through convective heat exchange. Larger beetles are surrounded by thicker boundary layers which reduce convective heat exchange with the air per unit surface area (Mitchell, 1976; Stevenson, 1985). When shape is held constant, increasing body size increases both the area intercepting solar radiation and the area available for heat loss, so surface area alone does not explain higher steady-state body temperature. Instead, larger beetles lose heat less efficiently per unit area for a given temperature difference with the environment, allowing absorbed solar energy to result in higher steady-state body temperatures. This difference was also apparent in the environments producing the highest predicted body temperatures. For the smallest beetles, the highest steady-state body temperatures occurred in arid north-western Western Australia, where air temperatures exceeded 40°C and the beetles remained relatively closely coupled to the surrounding air. Larger beetles, in contrast, could develop much greater temperature excesses at lower air temperatures when high solar radiation coincided with very low wind.

Our model represents an immobile, non-thermoregulating beetle exposed to a fixed set of environmental conditions with no metabolic heat production. The steady-state predictions therefore represent the potential effect of morphological traits if those conditions are maintained long enough for body temperature to equilibrate. The 60 s predictions instead describe the short-term thermal response after a beetle initially at air temperature is exposed to those same conditions. In the field, jewel beetles can move between microhabitats and alter their exposure to solar radiation and wind before steady state is reached, so realised body temperatures will also depend on behaviour and exposure duration. Flight would alter the heat budget further through increased convective exchange and metabolic heat production and was not considered here.

Body shape had smaller but consistent effects on steady-state body temperature during the daytime. Increasing relative width reduced steady-state body temperature whereas increasing relative height increased it. Increasing relative width results in a flatter, broader ellipsoid whereas increasing relative height results in a taller, narrower one. The posture used in the model was intermediate between parallel and perpendicular to the sun and averaged the minimum and maximum silhouette areas. Increasing relative width therefore reduced the projected area exposed to direct solar radiation relative to the total surface area available for heat exchange, whereas increasing relative height increased this ratio, producing lower and higher steady-state body temperatures, respectively (Kingsolver & Moffat, 1982; Mitchell, 1976; Ostwald et al., 2026). The negligible shape effects at night are consistent with the loss of direct shortwave heating, although body shape can still influence longwave radiative and convective heat exchange. Relative height also consistently increased time to equilibrium across all climatic conditions. At constant mass, increasing relative height produces a more compact ellipsoid by making all three axes more similar in length. Since more spherical bodies have a lower surface-area-to-volume ratio, this reduces the surface available for heat exchange relative to thermal mass, which increases heating and cooling time (Okie, 2013). By contrast, relative width had almost no effect on time to equilibrium.

Taken together, our results show that the thermal effect of solar absorptivity depends on the interaction between body size and the environment in which an insect experiences it. The potential magnitude of this effect is at most a few degrees Celsius, and the natural variation in body size of jewel beetles has an effect of similar magnitude. High absorptivity can therefore provide additional passive warming under cool, sunny conditions, while the same trait can increase thermal load when high solar radiation coincides with low wind.

Behaviour and exposure duration will further determine how much of this potential thermal effect is realised as insects move among microhabitats. The magnitude of the effect we predict is also constrained by the absorptivity range observed in jewel beetles, and other taxa or life stages spanning a wider range could show larger thermal differences.

## Data Availability Statement

The data and R code supporting this study are available on Figshare at https://doi.org/10.26188/34715004.

## Acknowledgements

We thank Katrina Rankin, Bryonny Margetts and Amanda Franklin for their assistance with fieldwork.

## Author contributions

Avichal Tatu, Devi Stuart-Fox, Michael Kearney, Nanfang Yu and Ann Roberts contributed to conceptualisation; Avichal Tatu, Devi Stuart-Fox and Michael Kearney designed the methodology; Avichal Tatu, Laura Ospina-Rozo and Nanfang Yu collected the data; Avichal Tatu analysed the data; Avichal Tatu, Devi Stuart-Fox and Michael Kearney led the writing of the manuscript. All authors reviewed and edited the manuscript, contributed critically to the drafts and gave final approval for publication.

