## Supporting Information for "How body size and microclimate determine when solar absorptivity matters for insect body temperature"

**Contents**

Methods S1. Solar absorptivity measurements and cross-dataset comparison

Methods S2. Mid-infrared emissivity measurements

Methods S3. Morphological measurements

Methods S4. Infrared thermography and environmental measurements

Table S1. Biophysical-model parameters and settings

Table S2. Field sampling and regional microclimate reconstruction used for validation

Table S3. Climate-block definitions and realised environmental conditions

Table S4. Model-validation statistics

Table S5. Representative cool-sunny and warm low-wind conditions used in the Discussion

Figure S1. Comparison of *Temognatha bruckii* reflectance spectra between datasets

Figure S2. Local effects of morphological traits on time to equilibrium

Figure S3. Environmental modulation of morphological-trait effects

Figure S4. Direct one-factor-at-a-time effects of environmental drivers

Figure S5. Spatial effect of solar absorptivity on steady-state body temperature under three wind-shelter scenarios

**Methods S1. Solar absorptivity measurements and cross-dataset comparison**

We used published reflectance data for 17 jewel beetle species from Wang et al. (2021) and added measurements from four species, with five individuals measured per species. *Temognatha bruckii* was represented in both datasets, allowing us to assess comparability between the two sets of measurements.

For the four newly measured species, total hemispherical reflectance was measured using an integrating sphere (Ocean Optics Inc.) equipped with a tungsten-halogen light source and a bifurcated fibre optic connected to USB2000+ and NIRQuest spectrophotometers (Ocean Optics Inc., Dunedin, FL, USA). The system was calibrated against 99% white and 2% black standards. Reflectance was measured from 300-1700 nm, following the wavelength range used to calculate broadband solar reflectivity in Wang et al. (2021).

For species with separate colour patches, measurements were taken separately from each region and weighted by the proportional area occupied by each patch, measured from photographs in ImageJ (Schneider et al., 2012), to calculate overall body reflectance. Broadband solar reflectivity was calculated by weighting spectral reflectance by the irradiance spectrum of the solar simulator and integrating across 300-1700 nm (Smith et al., 2016; Wang et al., 2021). Solar absorptivity (α) was then calculated as 1 - R, where R is broadband solar reflectivity expressed as a proportion.

For *T. bruckii*, the species common to both datasets, body reflectivity was 29.60% in our measurements compared with 28.36% in Wang et al. (2021), showing close agreement between datasets. The corresponding reflectance spectra are shown in Figure S1. Across the combined dataset, reflectivity ranged from 6.89 to 33.48%, corresponding to solar absorptivities of 0.67-0.93; the minimum and maximum observed absorptivity values were used as the empirical bounds for the sensitivity analyses.

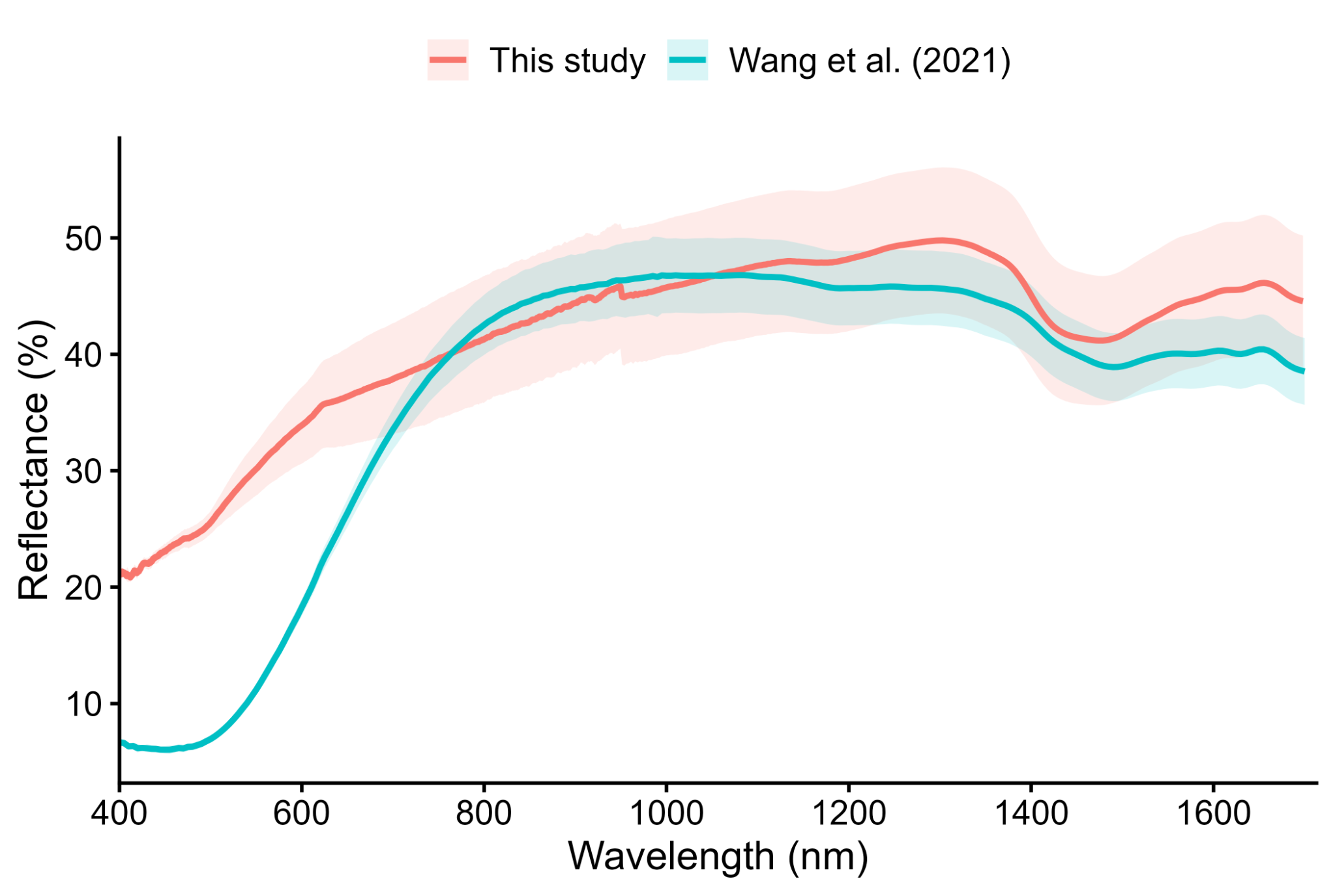

**Figure S1.** Comparison of *Temognatha bruckii* body reflectance spectra measured in this study and reported by Wang et al. (2021) across 400-1700 nm. The close agreement between datasets supports combining published and newly measured reflectance data when defining the empirical solar-absorptivity range.

**Methods S2. Mid-infrared emissivity measurements**

Mid-infrared emissivity was measured on five individuals per species across five species: *Castiarina octospilota*, *C. scalaris*, *C. sexplagiata*, *Temognatha bonvouloirii* and *T. bruckii*. Spectral reflectance and transmittance were measured using a Bruker Vertex 70v Fourier-transform infrared (FTIR) spectrometer.

Hemispherical reflectance was measured using a gold-coated integrating sphere calibrated against a gold reference standard. Transmittance was measured using a Bruker Hyperion 1000 microscope with reflective objectives and referenced to an open-beam measurement. Spectra were acquired over 3-16 µm and smoothed prior to analysis. Measurements were taken from the elytra on both dorsal and ventral surfaces, as well as from the wing, pronotum and prosternum, to assess whether transmittance differed between orientations.

ε(λ) = 1 - R(λ) - T(λ)

Spectral emissivity was calculated from reflectance and transmittance following Kirchhoff's law, as used previously to characterise mid-infrared emissivity in insect wings (Tsai et al., 2020). Broadband emissivity was estimated by weighting measured spectral emissivity by the Planck function at five temperatures spanning the biologically relevant range (293, 300, 312, 333 and 343 K). For each temperature, the weighted spectra were integrated over wavelength and normalised by the integral of the Planck function alone to obtain a broadband emissivity estimate. Emissivity was consistent across this temperature range and variation among individuals was small (0.88-0.90). The median broadband emissivity (ε = 0.90) was therefore used as a fixed model input in all subsequent analyses.

**Methods S3. Morphological measurements**

Morphological measurements of length, width, height and mass were taken from 90 individuals across 18 species selected to capture a broad range of body sizes and masses among Australian jewel beetles. Species ranged from small *Castiarina* approximately 7 mm in body length to large *Temognatha* approaching 45-50 mm, spanning more than two orders of magnitude in body mass. The morphological and absorptivity datasets only partially overlapped because species in each dataset were selected to capture variation in the traits of interest.

Maximum body length (a), width (b) and height (c) were measured using digital callipers (±0.01 mm). Relative width (b/a) and relative height (c/a) were calculated as descriptors of body proportions. This parameterisation allowed variation in body shape to be separated from overall body size when testing its effect on heat exchange. Body mass was measured using a Mettler Toledo XS205 DualRange analytical balance (repeatability <0.1 mg). In the model, mass determined body volume, while relative width and relative height determined how this volume was distributed among the three ellipsoid axes.

**Methods S4. Infrared thermography and environmental measurements**

Field validation was conducted on *Temognatha bonvouloirii* in Western Australia and *Castiarina sexplagiata* in Victoria. Beetle body temperature was measured using a FLIR T540 thermal camera fitted with a 24° lens (FLIR Systems, Wilsonville, OR, USA; 464 × 348 pixels). Thermal images were analysed in FLIR Thermal Studio.

For image processing, emissivity was set to 0.90 based on our beetle emissivity measurements. Reflected apparent temperature was set to ambient air temperature, and atmospheric temperature, object distance and related correction settings were specified from field measurements where available. Images were taken from approximately 1 m from the beetle, and this object distance was set in the camera at image capture. Where correction parameters were not recorded at image capture, they were applied during post-processing in FLIR Thermal Studio using the corresponding hourly environmental measurements. Body temperature was extracted from the dorsal surface.

Local air temperature, relative humidity and wind speed were measured using a Kestrel 5500 weather meter. Wind speed was averaged over 30-s sampling windows. Ground temperature and sky temperature were measured using an infrared thermometer. Measurements were taken at approximately hourly intervals and at a fixed height of approximately 1.5 m.

**Table S1.** Fixed biophysical-model parameters and settings used in the sensitivity and spatial analyses. Parameters not varied as focal traits were held constant so that model outputs reflected variation in the focal morphological traits and environmental conditions.

| **Parameter / setting** | **Value** | **Description** |
| --- | --- | --- |
| Posture | "a" | Intermediate posture |
| Substrate absorptivity | 0.30 | Fixed across analyses. |
| Longwave radiative temperature | Mean of sky and ground temperature | Used as the longwave radiative temperature input. |
| Beetle emissivity | 0.90 | Median measured broadband emissivity. |
| Evaporative heat loss | Not included | No evaporative cooling was included in the model. |
| Body specific heat | 3073 J kg⁻¹ K⁻¹ | Fixed across analyses. |
| Flesh thermal conductivity | 0.5 W m⁻¹ K⁻¹ | Fixed across analyses. |
| Initial body temperature for transient simulations | Air temperature | Used for calculation of body temperature after 60 s. |

**Table S2.** Field sampling and regional microclimate reconstruction used for model validation. The coordinates listed here are representative regional coordinates used to drive micro_silo; they are not the coordinates of every individual beetle observation.

| **Region** | **Focal species** | **Sampling period** | **Body-temperature n** | **Operative-temperature n** | **Representative micro_silo coordinate** | **Regional reconstruction settings** |
| --- | --- | --- | --- | --- | --- | --- |
| Victoria | *Castiarina sexplagiata* | Spring 2024 | 58 (50 with matched local inputs) | 58 (50 with matched local inputs) | 37.9678°S, 145.3987°E | Slope = 35°; aspect = 0°; wind factor = 0.5; microclima = 1 |
| Western Australia | *Temognatha bonvouloirii* | January 2025 | 31 | 30 | 29.5785°S, 117.0477°E | Slope = 0°; aspect = 0°; wind factor = 1.0; microclima = 0 |

**Table S3.** Climate-block definitions and realised environmental conditions for the 60 hourly environments sampled within each block. Daylight was defined as solar zenith angle < 90°, night as solar zenith angle >= 90°, and cool daytime conditions were restricted to hours at or before solar noon.

| **Climate block** | **Temperature criterion** | **Wind criterion** | **n** |
| --- | --- | --- | --- |
| Hot day, calm | TA >= 36.0°C (93rd percentile of daytime TA) | Wind ≤ 1.0 m s⁻¹ | 60 |
| Hot day, windy | TA >= 40.8°C (99th percentile of daytime TA) | Wind ≥ 3.5 m s⁻¹ (75th percentile of daytime wind) | 60 |
| Cool day, calm | TA <= 12.2°C (10th percentile; at or before solar noon) | Wind ≤ 1.0 m s⁻¹ | 60 |
| Cool day, windy | TA <= 12.2°C (10th percentile; at or before solar noon) | Wind ≥ 3.5 m s⁻¹ (75th percentile of daytime wind) | 60 |
| Cold night, calm | TA <= 6.2°C (10th percentile of nighttime TA) | Wind ≤ 0.7 m s⁻¹ (10th percentile of nighttime wind) | 60 |
| Cold night, windy | TA <= 6.2°C (10th percentile of nighttime TA) | Wind ≥ 2.6 m s⁻¹ (75th percentile of nighttime wind) | 60 |

**Realised climate summary (mean ± SD)**

| **Climate block** | **Air temp. (°C)** | **RH (%)** | **Wind (m s⁻¹)** | **Shortwave (W m⁻²)** | **Solar zenith (°)** | **Ground temp. (°C)** | **Sky temp. (°C)** |
| --- | --- | --- | --- | --- | --- | --- | --- |
| Hot day, calm | 37.5 ± 1.4 | 33.1 ± 12.3 | 0.87 ± 0.16 | 777 ± 148 | 26.2 ± 10.0 | 68.4 ± 11.7 | 26.2 ± 3.9 |
| Hot day, windy | 42.1 ± 0.9 | 17.8 ± 8.3 | 4.74 ± 0.96 | 903 ± 152 | 17.5 ± 10.0 | 64.0 ± 4.8 | 26.3 ± 4.7 |
| Cool day, calm | 6.8 ± 3.9 | 81.1 ± 21.8 | 0.79 ± 0.18 | 125 ± 118 | 80.8 ± 8.2 | 7.6 ± 6.3 | -12.3 ± 6.5 |
| Cool day, windy | 9.8 ± 1.9 | 60.8 ± 17.8 | 4.28 ± 0.88 | 181 ± 157 | 72.0 ± 16.3 | 10.9 ± 3.9 | -5.7 ± 7.0 |
| Cold night, calm | 3.1 ± 2.2 | 84.1 ± 16.3 | 0.54 ± 0.17 | 0 ± 2 | 90.0 ± 0.0 | 0.1 ± 2.5 | -16.4 ± 4.8 |
| Cold night, windy | 4.5 ± 1.7 | 62.6 ± 14.6 | 3.13 ± 0.49 | 0 ± 2 | 90.0 ± 0.0 | 2.5 ± 1.6 | -14.4 ± 5.7 |

Table S4. Validation of predicted steady-state body temperature against observed beetle body temperature and operative temperature using reconstructed microclimate inputs and primarily measured local environmental inputs. Pearson correlation (r), root mean square error (RMSE) and bias are reported for each comparison.

| **Region** | **Environmental inputs** | **Validation target** | **n** | **Pearson r** | **RMSE (°C)** | **Bias (°C)** |
| --- | --- | --- | --- | --- | --- | --- |
| Victoria | Reconstructed microclimate | Observed body temperature | 58 | 0.54 | 5.81 | -3.44 |
| Victoria | Reconstructed microclimate | Operative temperature | 58 | 0.53 | 7.48 | -4.91 |
| Victoria | Measured local inputs | Observed body temperature | 50 | 0.71 | 3.96 | 0.17 |
| Victoria | Measured local inputs | Operative temperature | 50 | 0.89 | 3.29 | -1.29 |
| Western Australia | Reconstructed microclimate | Observed body temperature | 31 | 0.85 | 3.34 | -1.35 |
| Western Australia | Reconstructed microclimate | Operative temperature | 30 | 0.88 | 2.92 | 0.14 |
| Western Australia | Measured local inputs | Observed body temperature | 31 | 0.91 | 2.40 | -0.55 |
| Western Australia | Measured local inputs | Operative temperature | 30 | 0.92 | 2.57 | 1.00 |

Table S5. Representative conditions illustrating how the thermal effect of solar absorptivity changes with body mass and microclimate. Low and high absorptivity correspond to the observed empirical bounds (α = 0.665 and 0.931).

| Condition | Region | Body-mass scenario | Lon, lat | Air temp. (°C) | Shortwave (W m⁻²) | Wind (m s⁻¹) | Tcf at low α (°C) | Tcf at high α (°C) | ΔTcf (°C) |
| --- | --- | --- | --- | --- | --- | --- | --- | --- | --- |
| Cool-sunny | Australian Alps / Southeastern Highlands | Minimum mass | 148.32, -36.60 | 7.6 | 570 | 0.20 | 13.66 | 16.10 | 2.44 |
| Cool-sunny | Australian Alps / Southeastern Highlands | Maximum mass | 148.32, -36.60 | 7.6 | 570 | 0.20 | 20.73 | 26.01 | 5.28 |
| Warm, low-wind hotspot | Longreach-Opalton region | Minimum mass | 142.92, -23.40 | 28.7 | 535 | 0.13 | 35.27 | 37.67 | 2.40 |
| Warm, low-wind hotspot | Longreach-Opalton region | Maximum mass | 144.12, -23.40 | 28.1 | 555 | 0.10 | 44.21 | 49.90 | 5.69 |

Note. The cool-sunny example represents a single realised pixel-hour and uses the same conditions for minimum and maximum body mass. The warm low-wind examples show the grid cells with the largest mean absorptivity effect across the ten locally highest-ranked shortwave radiation/(wind speed + 0.1) hours for each body mass. Environmental values for these rows are means across those selected hours.

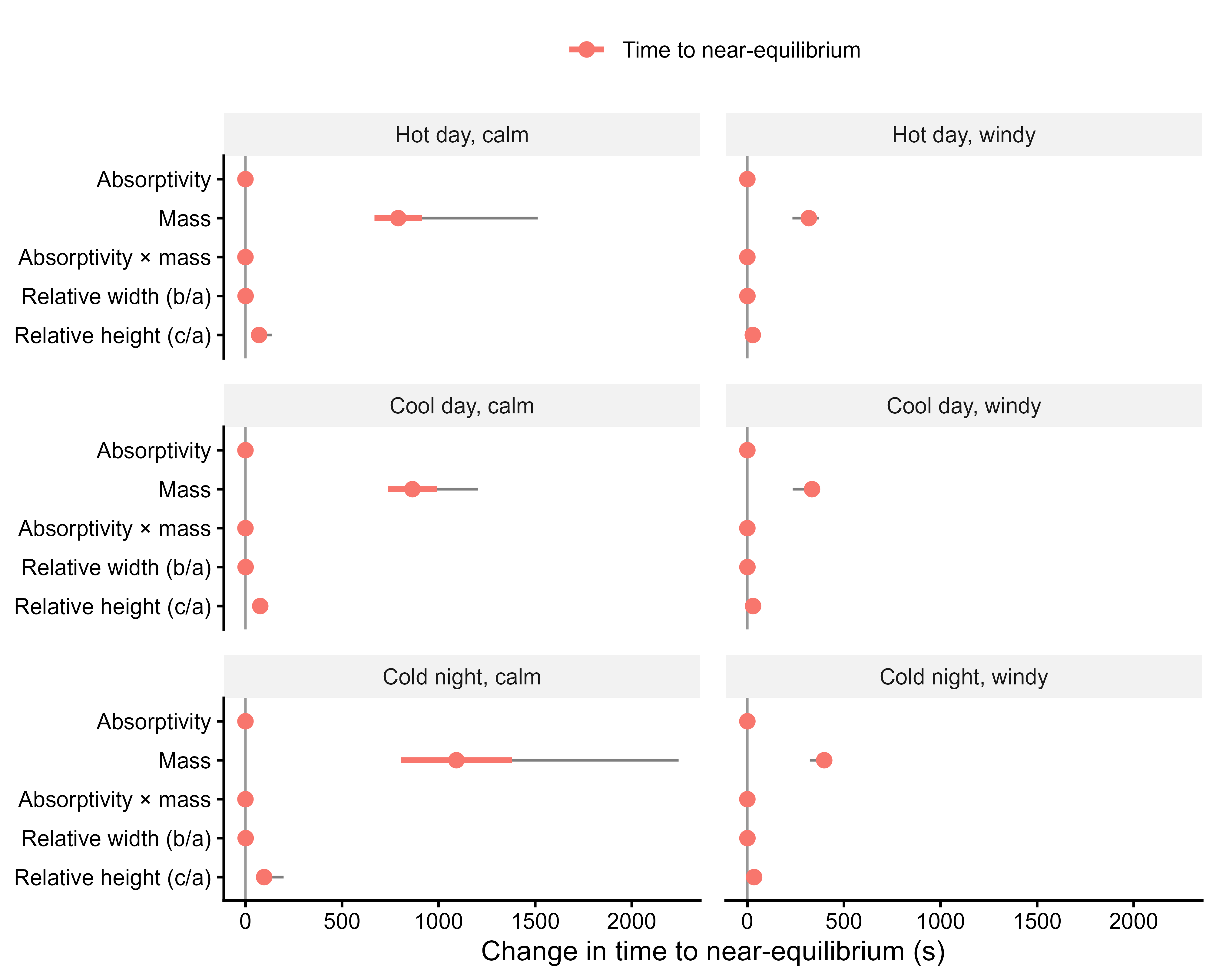

**Figure S2.** Local one-factor-at-a-time sensitivity of time to equilibrium to solar absorptivity, body mass, relative width, relative height and the absorptivity × mass interaction across six climate blocks. Points show the mean effect across 60 realised hourly environments within each block, thick bars show ±1 SD and thin whiskers show the minimum-maximum range. Trait effects are the difference in predicted time to equilibrium between the maximum and minimum observed value of each focal trait. Body mass dominated the response in every climate block, whereas absorptivity had no effect on equilibration time.

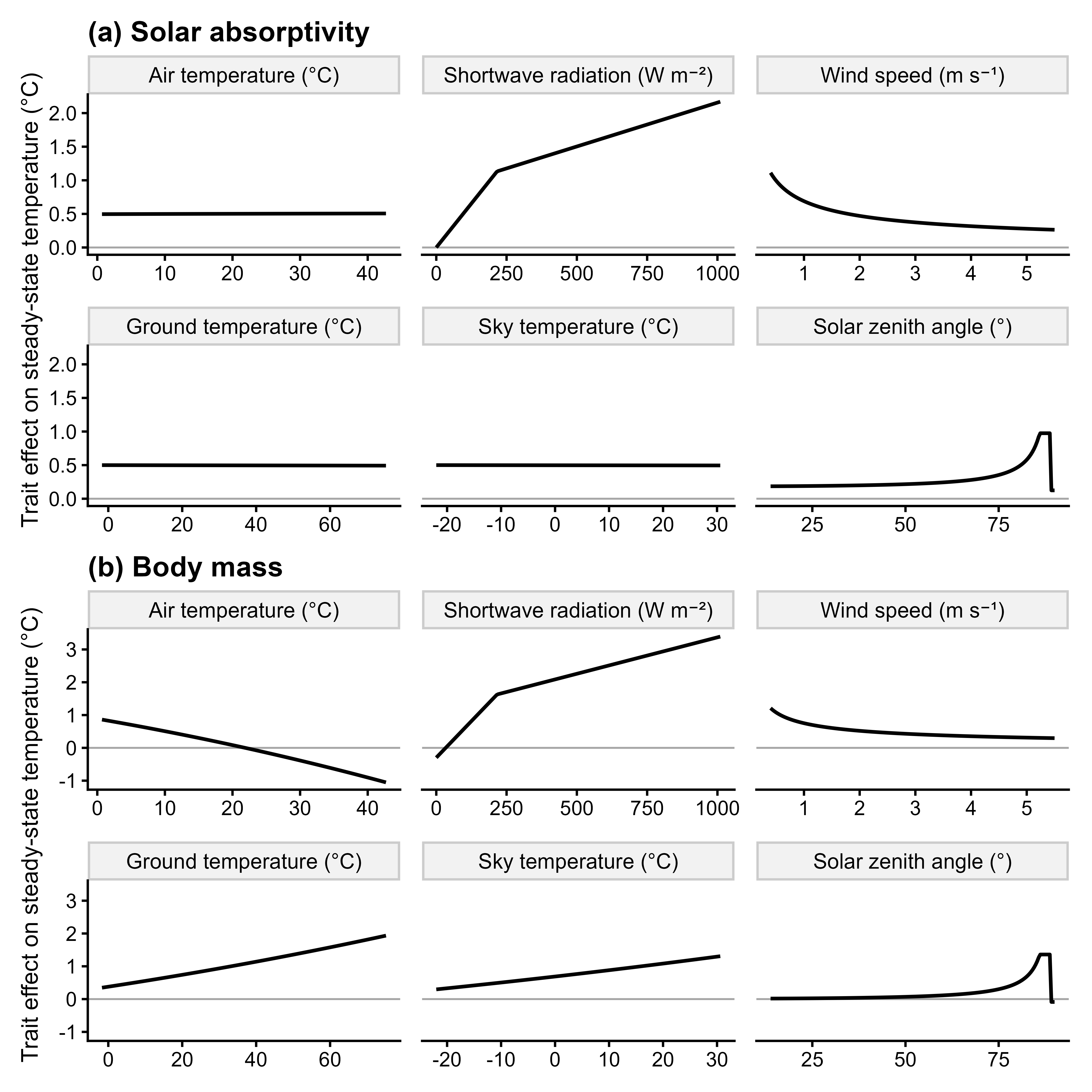

**Figure S3.** Environmental modulation of the steady-state thermal effects of (a) solar absorptivity and (b) body mass. Lines show the effect of each focal trait on steady-state body temperature ($T_{c,\mathrm{final}}$) as individual environmental drivers were varied across their 5th–95th percentile ranges. Trait effects were calculated as the difference in predicted $T_{c,final}$between the maximum and minimum observed value of the focal trait, while all other morphological traits were held at the midpoints of their observed ranges and all other environmental variables were held at their overall climate-panel medians. Environmental drivers were varied independently, so covariance among environmental variables was not retained in this analysis.

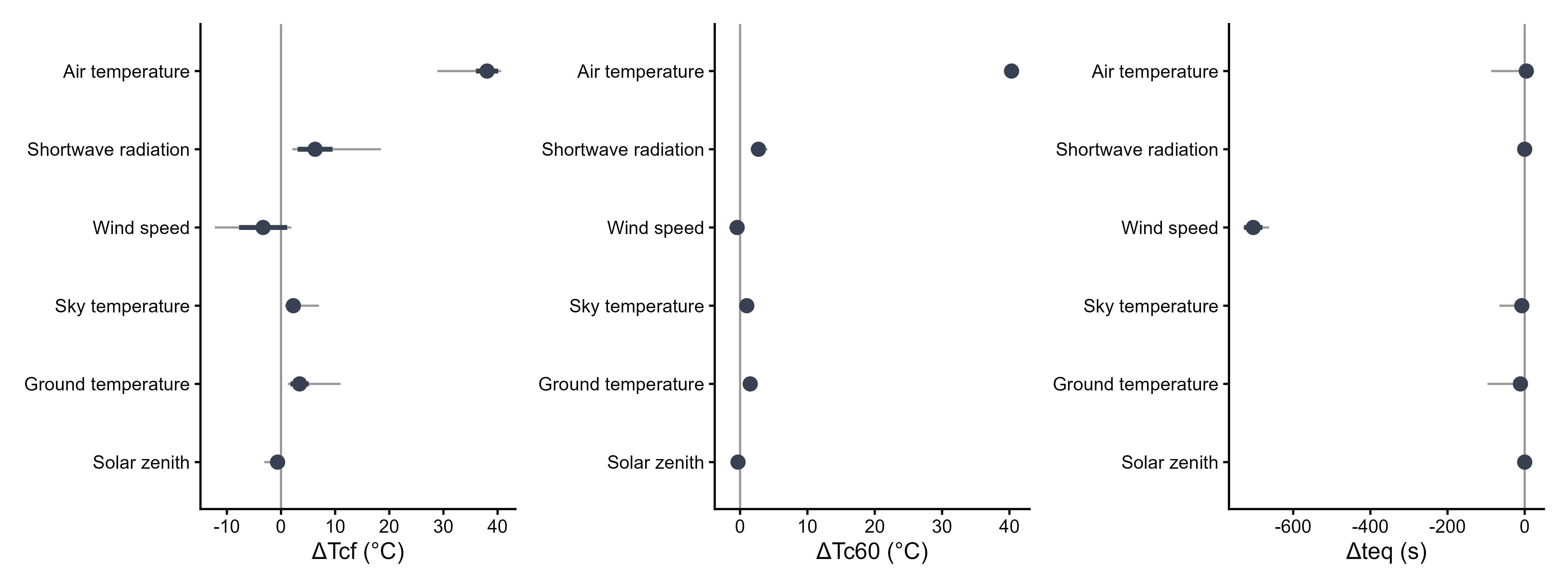

a)

c)

b)

**Figure S4.** Direct one-factor-at-a-time effects of environmental drivers on (a) steady-state body temperature, (b) body temperature after 60 s and (c) time to equilibrium. Each environmental driver varied across the global 5th-95th percentile range while the remaining environmental variables retained the values of each sampled environment. Points show mean effects across the 360 sampled environments, thick bars show ±1 SD and thin whiskers show the minimum-maximum range.

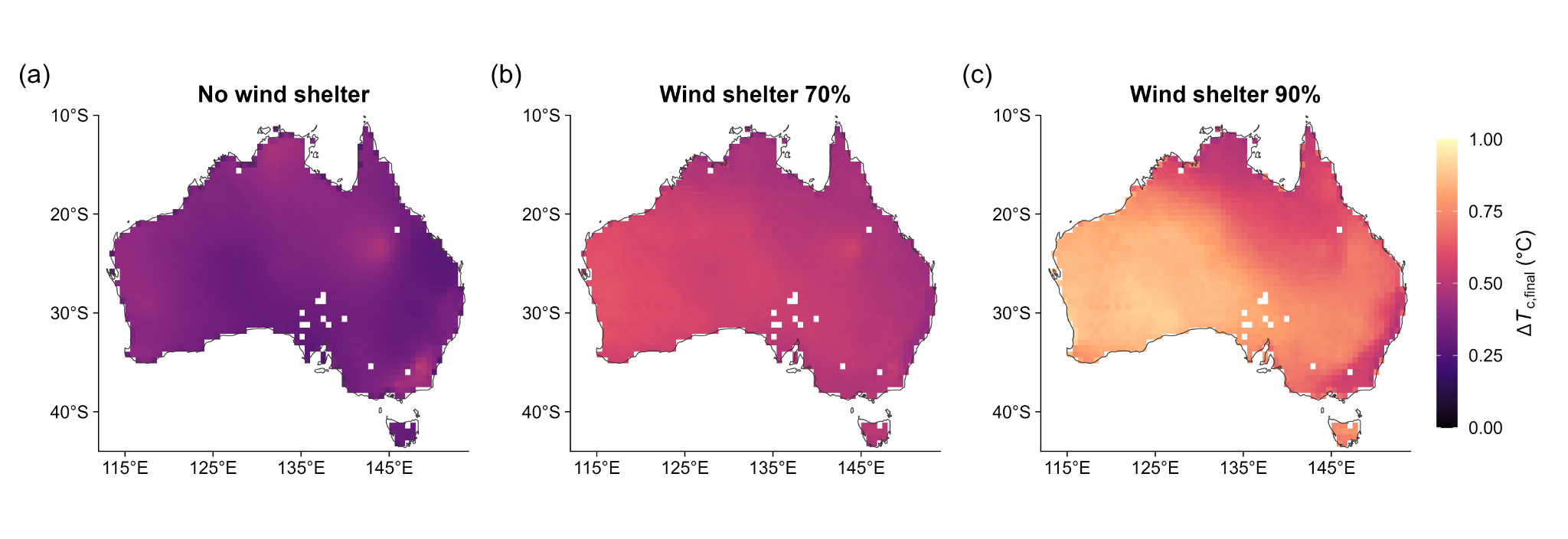

**Figure S5**. Spatial variation in the effect of solar absorptivity on predicted steady-state body temperature (ΔT*_c,final_*) across Australia during daylight hours in October–December 2017. Maps show mean ΔT*_c,final_* when wind speed was (a) left unchanged, (b) reduced by 70% whenever wind speed exceeded 1.5 m s⁻¹, and (c) reduced by 90% whenever wind speed exceeded 1.5 m s⁻¹. All other environmental variables and morphological traits were held unchanged. Warmer colours indicate a larger predicted thermal effect of the observed absorptivity range.
